# Quantifying the effect of metal interactions on growth rate in *Saccharomyces cerevisiae*

**DOI:** 10.1101/2025.09.18.675698

**Authors:** Penelope C. Kahn, Sarah P. Otto

## Abstract

Environmental stressors often co-occur, yet their combined effects on organisms remain poorly understood. As multifactorial anthropogenic changes intensify and alter environmental selection pressures, our understanding of stressor interactions is becoming increasingly important. However, established methods for characterizing the effects of stressor interactions can involve considerable experimental and computational labour. Here, we investigate how pairwise combinations of six divalent metal ions (Cd^2+^, Co^2+^, Cu^2+^, Mn^2+^, Ni^2+^, and Zn^2+^) affect the growth of *Saccharomyces cerevisiae*, using a high-throughput assay to generate concentration-response surfaces for all 15 combinations. We introduce *δ*, an easily calculable metric that quantifies the toxicity of a mixture relative to the toxicities of its individual components, and compare it to an established metric for synergism/antagonism, *a*, determined by calculating deviation from a null model of additivity, i.e., “concentration addition”. Yeast growth rates reveal that metal mixture toxicity varies widely from the additive expectation with the effect of the combination ranging from greatly enhanced toxicity to greatly attenuated toxicity. Combinations with copper were more toxic than expected, and combinations with manganese or nickel were less. These trends correspond to known redox activities and metal-binding properties, pointing to possible mechanistic underpinnings rooted in oxidative stress and metal cofactor displacement. Furthermore, we find that *a* correlates with overlap in known metal resistance genes and similarity in redox potential, offering predictive insight into effects of metal interactions. While *a* provides a model-based measure of synergism or antagonism, *δ* serves as an intuitive, concentration-robust descriptor of ecological impact. Together, these metrics highlight the complexity of metal-metal interactions and the importance of accounting for nonadditive effects in ecotoxicological assessments. This framework provides a basis for evaluating mixture toxicity across diverse taxa and stressor types, with implications for both evolutionary biology and environmental risk assessment.

## Introduction

Severe environmental change requires that species adapt or go extinct, but the severity of the challenge is not predictable based on the effects of individual stressors in isolation. As human population size and technological innovation have grown exponentially over the last century, so too have the volume and diversity of our waste products. Organisms inhabiting polluted environments are thus subject to combinations of stressors, the components of which can interact to modify the toxicity of the environment. Generally, hazardous waste must be consolidated and stored with like substances until it can be processed, but containment methods are susceptible to failure^1^. For example, failures in mine tailing storage since 1915 have caused land area destruction up to 5,000 km^2^, an estimated human death toll of 2,650, and untold declines in wildlife^2^. Mine tailings contain multiple toxic metals, creating conditions where organisms must tolerate complex mixtures of multiple toxic components. Though we may understand the biological effects of each component in isolation, it is challenging to predict their effects when combined^3,4,5^.

Efforts to model the effects of combined toxins began a century ago^6^ and have been developed and applied in pharmaceutical, agricultural, and ecotoxico-logical sciences^7^. The aim of this literature is to determine if interactions between components in a mixture either enhance or attenuate their isolated harms. “Synergism” occurs when the combined effect of two toxins is more harmful than expected from their individual effects, whereas “antagonism” occurs when the combined effect is milder than expected, implying some form of mitigation or interference between toxins.

Assessing interactions thus requires a null model for comparison. Here we focus on the null model of “concentration addition” (CA; also referred to as “method of isoboles” or “isobologram approach”^8^). CA assumes that, if two toxins were identical, their combined effect should be equivalent to doubling the concentration of either one. If an observed response to a mixture of toxins deviates from the null model predictions, an interaction can be inferred, with enhanced toxicity implying synergism and attenuation implying antagonism^9^. Because combining metals should satisfy the CA assumption that metals are similar in action (further described in the next section), we focus on this null hypothesis and use a method introduced by Haas et al.^10^ and further developed by Jonker et al.^9^ to test for significant deviations from the CA model using maximum likelihood methods.

Many of the first-row transition metals (e.g., Mn, Fe, Co, Ni, Cu, and Zn) are required in small amounts as micronutrients in the cell^11^. Their ionic abilities to donate and receive electrons, stabilize charged substrates, and selectively bind ligands make them useful cofactors in many enzymes, but these same properties become detrimental to cellular functioning when concentrations reach high levels^12^. Oxidative stress is a major consequence of excess metal ions, caused either directly by production of reactive oxygen species (ROS)^13^ or indirectly by depletion of antioxidants (e.g., the glutathione pool)^14^. Excess metal ions can also inactivate enzymes by displacing native metal cofactors^15^ or by binding to -SH groups in cysteine residues, destabilizing protein structure and inhibiting proper folding^16^. Some toxic properties specific to certain metals include direct DNA damage and prevention of DNA repair^17,18^, mitochondrial damage and dysfunction^19,20^, and destabilization of membranes directly through lipid peroxidation or indirectly through alteration of membrane potential^21,22,23^. While there has been significant effort to understand the cellular effects of many individual metals^16,24,25^, we know little about their combined effects at the molecular level, which is surprising given the abundant evidence that metal interactions can significantly impact environmental toxicity.

By contrast to sparse molecular investigations of metal mixture toxicity, numerous studies have examined the effects on an organismal or population level, investigating the synergism/antagonism of metal combinations in aquatic or soil invertebrates^26,27,28,29,30,31,32,33,34^, model or crop plants^35,36,37,38,39^, phytoplankton^40,41^, bacteria^35,42,43,44,45,46^, and various model vertebrate species^47,48,49,50^. While this constitutes a wide range of taxa, fungi are notably absent, and microbial species are significantly underrepresented given their biodiversity and myriad ecological roles from geochemical cycling to decomposition^51^. Furthermore, these studies frequently test only a small number of metal combinations, most often involving cadmium, copper, and zinc (see Discussion for a review of this literature).

Here we offer the first look at metal mixture toxicity in the model organism *Saccharomyces cerevisiae* using 15 pairwise combinations of six heavy metals (five essential elements: cobalt, copper, manganese, nickel, zinc; and one non-essential element: cadmium). Metals were first analysed on their own to determine their dose-response curves in isolation, which allowed us to choose concentrations with similar toxicity levels for the combinations. We then compared growth rates of an isogenic yeast strain in each single metal and their combination to observe the change in toxicity resulting from simultaneous exposure. We demonstrate our new measure of cumulative toxicity, *δ* (described in the next section), and compare it to the interaction term, *a*, showing that pairs of metals interact along a spectrum ranging from strongly antagonistic to strongly synergistic, with twelve of fifteen metal pairs deviating significantly from CA expectations. We also find that *δ* provides a fairly robust measure of the reduction in fitness caused by combining two metals that remains nearly constant for a given metal pair across a concentration gradient of increasing severity. Our findings highlight the complexity of metal-metal interactions and their implications for ecotoxicology and evolutionary adaptation. In natural environments, organisms are rarely exposed to single toxicants, and understanding the interactions between stressors is essential for predicting evolutionary responses and risk of extinction in the face of severe environmental pollution.

### Measures of mixture toxicity

We used two methods to quantify the toxic impact of combining environmental stressors (referred to as “toxins” generically and “metals” when discussing the experiments). Our first measure, *δ*, describes how harmful the stressor combination is relative to the individual stressors, given a certain degree of harm caused by each of the two stressors in isolation. This intuitive and easily calculable measure is ecologically relevant when the researcher’s goal is to assess the danger an organism faces when encountering the cumulative effect of two stressors at once. Here we use the term “cumulative” to describe the combined effect of multiple stressors without implying that their effects combine additively. Specifically, we define a “cumulative toxicity” measure, *δ*, as:

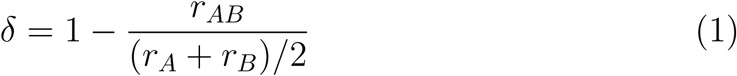

where *r_AB_*is the maximum growth rate observed in the combination of toxins, which is measured relative to the average growth in the single toxins (at the same concentrations as in the mixture), (*r_A_* + *r_B_*)*/*2 (see Figure 1B).

**Figure 1:**
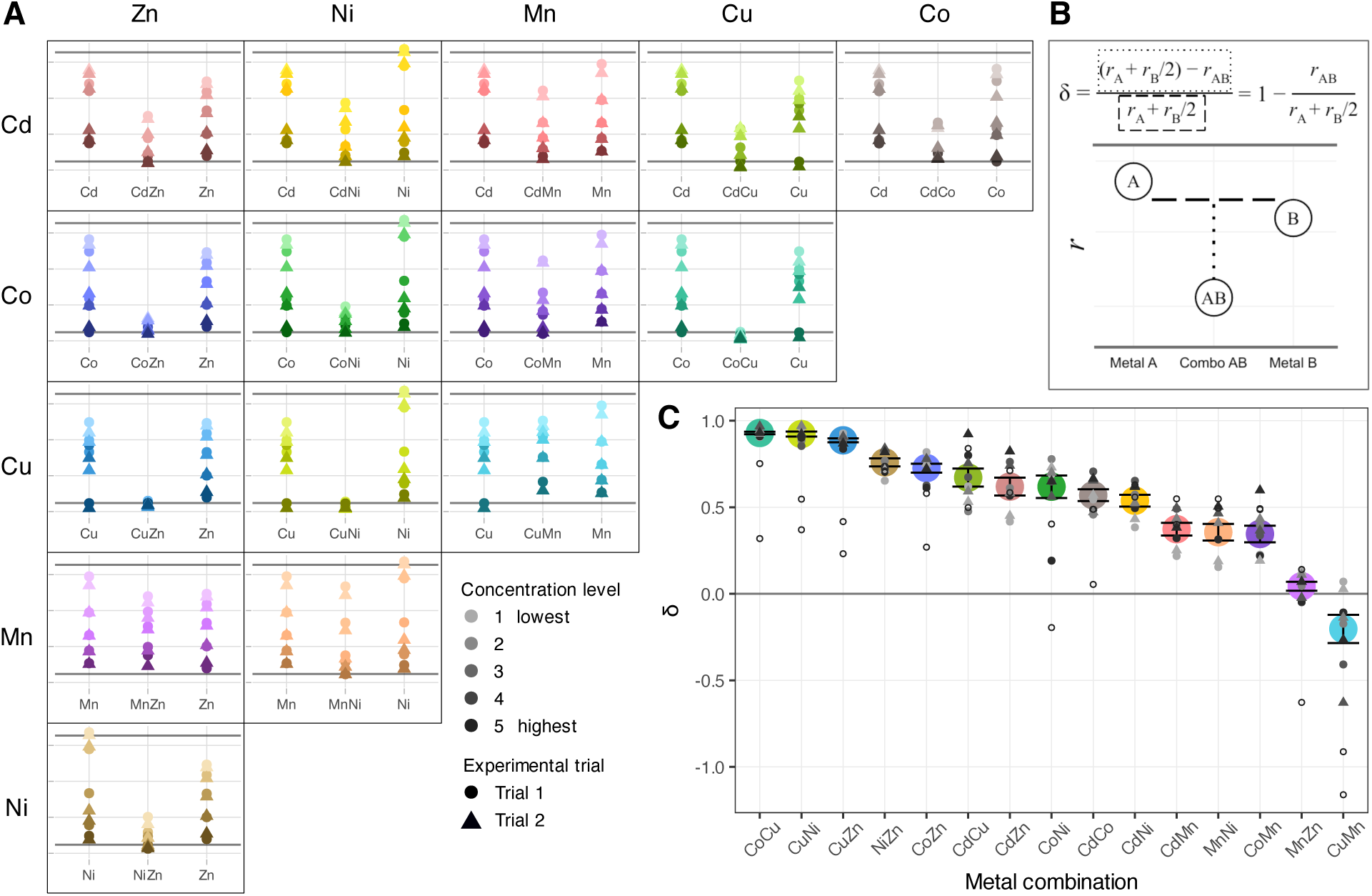
Measurement and quantification of metal mixture toxicity on *S. cerevisiae* growth rate. (A) Maximum growth rates (*r*; y-axis) are derived from growth curves of a single *S. cerevisiae* genotype exposed to six metals (Cd, Co, Cu, Mn, Ni, Zn) and their pairwise combinations (holding constant the single metal concentrations). Growth rates in single metals are displayed on the left and right of growth rates in their combination (middle). Horizontal bars on top and bottom correspond to *r* of yeast in YPAD (positive control) and water in YPAD (negative control). Color intensity corresponds to the five target concentrations, which aimed to reduce maximum growth rate in the single metals from ∼0.3 to 0.25, 0.2, 0.15, 0.1, 0.05 (light to dark). Points represent mean from three technical replicates, with circles and triangles indicating data from one of the two independent trials (see Table S1 for metal concentrations, which were adjusted slightly after the first trial). (B) Visual representation of *δ* calculated by taking the quotient of the difference in maximum growth rate in single and combined (“combo”) metals divided by the average growth in single metals at roughly equitoxic concentrations. (C) *δ* values for each metal combination calculated from data in panel A. Large colored dot indicates average *δ* across trials and concentration levels 1-4 within a given metal combination (± SE). Small grey points represent data from concentrations levels 1-4; open circles represent data from concentration level 5, where growth was negligible, which were excluded from statistical analyses of *δ*. Color intensity and shape have the same meaning as in panel A.

Importantly, the degree of toxicity of each stressor was held roughly constant (i.e., *r_A_*∼ *r_B_*) when assessing the impact of the combinations at various concentrations, allowing us to measure *δ* over a gradient of stress levels, from mild to severe (“stress level” referring to a concentration that reduces fitness, here measured as maximum growth rate, by a given amount). This measure is scale free in the sense that multiplying all growth rates by any quantity (e.g., changing time units) does not alter *δ*. Empirically, as we show here using experiments with yeast, *δ* is nearly constant for a pair of metals over the gradient of stress levels applied (see Figure 1C). By contrast, *δ* was very sensitive to the identity of the metals combined. For some metal combinations, the cumulative effect of both was no more toxic than each metal on its own (*δ* near 0), while in other cases the cumulative effect was much more severe and did not allow any growth (*δ* near 1). Thus, the cumulative toxicity measure, *δ*, provides information about which combinations of toxins are particularly harmful and which are not.

As a second measure, we estimated the interaction value, *a*, of Haas et al.^10^. This quantity measures the extent to which a combination of toxins behaves differently from simply increasing the concentration of either toxin (the CA null hypothesis). In the determination of *a*, the toxicity of the combination is examined in reference to a null model, the choice of which requires assumptions to be made about the components’ mechanisms of action, whereas in the determination of *δ*, the toxicity of a mixture is examined relative to the toxicity of its components, which only requires observations of growth rates in each environment.

Following Haas et al.^10^, we estimate *a* by first assessing the toxicity of each stressor on its own, measuring the maximum growth rate across a concentration gradient for toxin *i*, yielding a dose-response curve that is then fit to a logistic function:

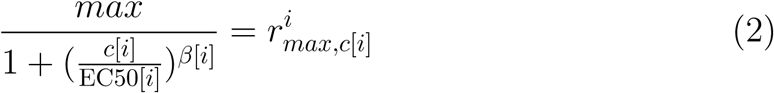

In equation 2, *c*[*i*] is the concentration of toxin *i*, 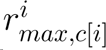 is the maximum growth rate observed at that concentration, max is the maximum growth rate in the absence of the toxin, EC50[*i*] is the concentration of toxin *i* that causes the growth to decline to half of this maximum value, and *β*[*i*] is proportional to the slope of the logistic function at the EC50[*i*] concentration.

According to Haas et al.^10^ and Jonker et al.^9^, the interaction between two toxins can then be measured by solving the following for *a*:

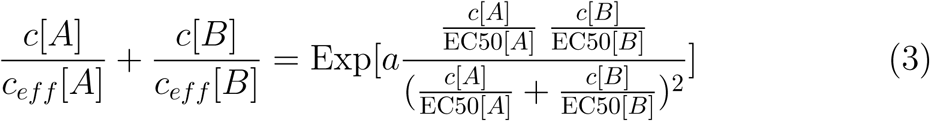

On the left-hand side of equation 3, the concentration of metal *i* used in the mixture, *c*[*i*], is divided by the effective concentration of the single metal *i*, *c_eff_* [*i*], that would be needed to observe the same growth rate as that seen in the mixture. Specifically, *c_eff_* [*i*] is estimated by solving equation 2 for the concentration of metal i that would yield the observed growth rate in the metal combination (*r_AB_*). On the right-hand side of equation 3, *a* measures the departure from the null hypothesis of concentration addition, while the fraction measures the evenness of the two toxins being combined (ranging from 0 when one substance is at much lower concentration than the other to a maximum of 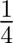 when both toxins are at equal concentrations, when measured relative to their EC50s).

If the two metals in the mixture were actually the same metal, then combining the two at equal concentrations must be equivalent to doubling either metal, so that *c_eff_* [*i*] = 2 *c*[*i*]. In this case, the left-hand side of equation 3 equals one, *a* equals zero, and the concentration addition hypothesis is satisfied. If, however, the metals are more toxic when combined, the effective concentration of each metal must be larger to account for the low growth in the mixture, leading to a negative estimate of *a* (“synergistic toxicity”). Conversely, if metals are less toxic when combined than expected, *a* becomes positive (“antagonistic toxicity”).

We note that evolutionary biologists often use a different null model when considering interactions, assuming that two stressors act multiplicatively on fitness (i.e., additively on growth rates). Epistasis, for example, is measured relative to this null, to describe how the fitness effect of a mutation depends on other alleles present in the genetic background. When considering mixture toxicity, the analogous model is called “independent action” (IA). IA assumes that each stressor acts independently on fitness and is relevant, for example, when stressors affect survival rates at different stages^52^. However, the effects of multiple metals are not expected to remain independent of each other in simultaneous exposure due to overlap in cellular uptake, response, and detoxification systems^53^. In fact, when combining two metals, each of which has a logistic effect on toxicity, IA is expected to underestimate how toxic the combination will be whenever adding two metals shifts the system from below to above the EC50.

Furthermore, as shown in Figure S1, *δ* values determined analytically from the CA model are stable over a range of concentrations, as long as the two metals provide similar stressor intensities (i.e., induce similar growth rates), which aligns with our empirical observations (Figure 1C). By contrast, the IA model does not predict a constant *δ* over a range of stressor intensities. We thus do not discuss the IA model further.

## Material and Methods

### Strains and Media

The source of yeast used in all experiments in study was a frozen sample of the ancestral haploid line “OLY077” described in^54^. Briefly, the original OLY077 stock was created by propagating an isogenic colony sample of a MAT-*α* haploid line, generated by crossing and sporulating two haploid lines of the W303 background from McDonald et al.^55^. We bottlenecked the OLY077 stock to a single colony again to minimize genetic variation in our experimental populations. Specifically, a frozen sample of the original OLY077 stock was inoculated in liquid rich medium (YPD (RPI Research Products Y20090) + 20 mg/L adenine hemisulfate, referred to as YPAD) and grown overnight at 30 ^◦^C. The saturated culture was diluted 1:1000, plated on YPAD + agar, and incubated at 30 ^◦^C for two days. A single colony was isolated and propagated in liquid YPAD overnight, shaking at 30 ^◦^C. A freezer sample, to be used as the source population for all experiments in this study, was prepared by mixing the saturated culture with 30% glycerol at a 1:1 ratio and stored at −80 ^◦^C.

Stock metal solutions (0.1 M) were prepared using the following metal salts in powder form: Cadmium sulfate, anhydrous, ACS, 99+% (Alfa Aesar 11861); Cobalt(II) sulfate, heptahydrate, 98% (Alfa Aesar A16201); Copper(II) sulfate, pentahydrate, 98+% (Sigma-Aldrich C3036); Manganese(II) chloride, tetrahydrate, 99+% (Sigma-Aldrich M3634); Nickel(II) sulfate, hexahydrate, ACS, 98% (Alfa Aesar 36336); Zinc sulfate, heptahydrate (Sigma-Aldrich Z0251). Measured powder was added to distilled-deionized, ultra-pure water (Barnstead B-pure and Barnstead Nanopure Diamond) and stirred until crystals dissolved. The solutions were autoclaved at 121 ^◦^C and 15 psi with a sterilization time of 20 minutes

### Growth rate assays

Growth trajectories were measured using a Bioscreen C, a microbiology workstation that automatically records turbidity of cultures in 100-well patented honeycomb plates at regular intervals using wide band filters (Oy Growth Curves Ab Ltd., 2009).

For each Bioscreen assay, we prepared media for three technical replicates of each condition in 1.5 mL microcentrifuge tubes by mixing aqueous metal stock solutions into YPAD (see Table S1 for concentrations), after which a volume of overnight yeast culture was added at a dilution of 1:100 (approximately 100,000 cells founding each well^56^). To ensure the negative controls were matched to the treatment conditions, blank samples were diverted from the treatment tube before adding yeast. Additional positive and negative controls were yeast in YPAD and water in YPAD, respectively. 120 *µ*L aliquots from the prepared tubes were loaded into wells in a randomized order across four 100-well plates, and experiments were run concurrently in two Bioscreen machines holding two plates each. Plates were incubated at 30 ^◦^C and shaken continuously at medium amplitude, and optical density (OD) was measured every 15 minutes using a wide-band (420-580 nm) filter.

Maximum growth rate (*r*), a common measure of fitness for microbial populations, was the response variable used to assess toxicity. All statistical analyses and data visualizations were performed in R v4.4.2^57^, unless otherwise stated. We conducted a non-parametric loess fit to the log of OD over time (using loess in R with degree=1 and span=0.1), the slope of which provides an estimate of the instantaneous growth rate. The maximum growth rate was then obtained by finding the maximum slope of a non-parametric loess fit to the log of OD over time (see details in Gerstein et al.^58^).

### Dose-response curves

We measured maximum growth rates across a gradient of concentrations for each metal (see Figure S3A for dose-response curves). Metal concentrations were chosen to yield a range of growth rates from normal growth (approximately 0.32 OD unit/hour in a pilot experiment as measured in YPAD with no added metal) to no growth over the 48-hour measurement period. A broad range of concentrations was tested first to identify the approximate EC50. Subsequent trials were performed, narrowing in on concentrations near the EC50 to increase resolution of the dose-response curve, used to inform the choice of concentrations used in the combined metal growth assays (Table S1).

### Growth in combined vs. single metals

The dose-response curves were used to determine the experimental concentrations for each single metal at each “concentration level”. Here, concentration level refers to concentrations of different metals that induce the same maximum growth rate (*r*). Concentration levels were chosen to span the range from normal to zero growth (target *r* for conc 1: 0.25; conc 2: 0.20; conc 3: 0.15; conc 4: 0.10; conc 5: 0.05; see horizontal lines in Figure S3B-G). Overall, the growth rates at the chosen concentrations aligned well to the targets. Significant deviations from target growth rates were assessed using a linear mixed-effects model (lmer) with concentration level and metal as fixed effects and trial as a random effect. Estimated marginal means were computed with emmeans, and identity contrasts tested whether deviations differed significantly from zero. Deviations from target *r* (standardized to zero across concentration levels) are displayed in Figure S3H with asterisks denoting a conservative confidence level of 0.01. Significant deviations include manganese and nickel in concentration level 1, cadmium and nickel in concentration level 2, cadmium in concentration level 3, copper in concentration level 4, and cadmium and copper in concentration level 5. Regardless, no deviation exceeded one-third the range of possible growth rates, so we consider differences in toxicities within a concentration level to be reasonable, and we include all maximum growth rate data (displayed in Figure 1A) in calculations of *δ* and *a*.

Within a given concentration level, the same mM concentrations for each individual metal were used across the single metal condition and all its double metal pairs (i.e., the mM concentration of cobalt in Co level 1 is identical to that of cobalt in CdCo level 1, CoCu level 1, etc.).

Metal-media stock solutions were prepared at twice the desired concentration. For single metal-media treatment solution, part of the 2X stock solution was taken and diluted by a factor of two in YPAD to obtain the desired concentration. For double metal-media treatment solutions, 2X stock solutions of the two metals were mixed in a 1:1 ratio, reaching the desired concentrations simultaneously.

### Quantifying metal interactions

Cumulative toxicity, *δ*, was calculated from equation 1 at each of the five concentration levels in both trials. Replicate estimates of *δ* across concentration levels were generally consistent, except for the most severe metal concentrations (conc 5), where growth was negligible leading to noisy estimates of *δ* (Figure S5). We thus include conc 5 in Figure 1 and Figure S3 and in the fitted surfaces but exclude them from statistical analyses of *δ*.

Using the same data, the interaction term *a* ^9^ was estimated by maximum likelihood in *Mathematica* (v14.2.0.0). In brief, the parameters of the model {*a*, *max*, EC50[1], EC50[2], *β*[1], *β*[2]} allow estimation of the maximum growth rate*r*for any given combination of metal concentrations, *c*[1] and *c*[2]. Equation 2 was used to solve for the effective concentrations of each single metal, *c_eff_* [1] and *c_eff_* [2], that would yield a maximum growth rate of *r*. Plugging these relationships and parameter values into equation 3 results in an equation that can be solved numerically for *r*. The likelihood of the observed growth rate data at *c*[1] and *c*[2], using both experimental trials (Table S1), was modeled as a normal deviate, with a mean of *r* and a constant error variance (*σ*^2^). We then maximized the sum of the log-likelihood (*lnL*) over all growth rates estimated for a pair of metals (including the single metals and the combination) to obtain maximum likelihood estimates of the parameters, given the growth rate data. A likelihood ratio test was conducted to assess support for the unconstrained model (allowing any value of *a*), compared to the null hypothesis of concentration addition (*a*=0), comparing 2 (*lnL_a_* − *lnL_a_*_=0_) to a *χ*^2^ distribution. To assess robustness of this analysis to the response measure used, we also calculated *a* using the difference between final and initial OD (*a*[OD]) instead of maximum growth rates (*a*[*r*]).

## Results

### Growth in combined vs. single metals

To assess the toxicity of metal combinations across stressor intensities, we calculated maximum growth rates of yeast grown in five concentrations of six metals (Cd, Co, Cu, Mn, Ni, and Zn) and all of their 15 pairwise combinations (Figure 1A). The presence of two metals in the same environment had varied effects on yeast growth, depending on the identity of the metals in the pair. As shown in Figure 1A, some combinations of metal, particularly CoCu and CuZn, did not allow growth even at what would be low, permissive concentrations of each metal in isolation, causing deep troughs when plotting growth rate for the metal combinations. For other combinations, notably MnZn and CuMn, however, yeast grew equally well, or even better when exposed to a combination of metals compared to growth in each individual metal, leading to relatively flat comparisons in Figure 1A. Growth in the remaining combinations fell between these examples (Figure 1A).

### Measures of ***δ***

To quantify the impact of combining metals on fitness, *δ* values were calculated from two independent experiments. The mean *δ* for each pair of metals ranged from −0.63 to +0.97 (Table 2, Figure 1C), with a left-skewed distribution (mean = +0.54, median = +0.61). *δ* remained consistent across concentration levels for most metal combinations, as indicated by low sample variances (mean *s*^2^ = 0.01), suggesting stable interaction effects across stressor intensities (Figure S6).

**Table 1:** Description of parameters.

| Parameter | Description | Value | Meaning |
| --- | --- | --- | --- |
| $r$ | Maximum growth rate; inflection point of loess-fit growth curve | $\sim 0.32$ | Normal growth |
| | | $\sim 0.02$ | No growth |
| $\delta$ | Cumulative toxicity; change in toxicity when combining stressors | $> 0$ | More toxic when together |
| | | $< 0$ | Less toxic when together |
| EC50 | Concentration that results in 50% reduction of growth rate from normal | EC50<sub>Co</sub> | 0.92 mM Co ($r \approx 0.16$) |
| | | EC50<sub>Zn</sub> | 4.87 mM Zn ($r \approx 0.16$) |
| $a$ | Interaction term determined by deviation from CA | $> 0$ | Synergistic toxicity |
| | | $< 0$ | Antagonistic toxicity |
| $\beta$ | Slope of an individual concentration-response curve | $\beta_{Ni} = 11.6$<br>$\beta_{Zn} = 4.6$ | Affects the width of the range of response (from 0 to 100% maximum effect) along a concentration gradient; Ni has a narrow window of response (large slope), Zn has a broad window of response (small slope) |
| $\delta^*$ | $\delta$ inferred from 3D surface fitted to data using $a$ | $-\delta^* > -\delta$ | Model underestimates benefit from combination |
| | | $-\delta^* < -\delta$ | Model overestimates benefit from combination |
| | | $+\delta^* > +\delta$ | Model overestimates harm from combination |
| | | $+\delta^* < +\delta$ | Model underestimates harm from combination |

**Table 2:** Numerical values of interaction terms.

| | Metal combination | Observed $\delta^\dagger$ | Inferred $\delta^{*\dagger}$ | $a^\ddagger$ | p-value |
| --- | --- | --- | --- | --- | --- |
| | CdCo | $0.55 \pm 0.06$ | $0.59 \pm 0.07$ | -0.16 | (n.s.) |
| | CdCu | $0.64 \pm 0.07$ | $0.66 \pm 0.05$ | -1.59 | ** |
| | CdMn | $0.36 \pm 0.05$ | $0.41 \pm 0.07$ | 1.25 | *** |
| | CdNi | $0.53 \pm 0.06$ | $0.57 \pm 0.08$ | 0.87 | *** |
| | CdZn | $0.59 \pm 0.07$ | $0.63 \pm 0.07$ | 0.20 | (n.s.) |
| | CoCu | $0.92 \pm 0.01$ | $0.99 \pm 0.01$ | -2.41 | *** |
| | CoMn | $0.29 \pm 0.04$ | $0.38 \pm 0.06$ | 1.93 | *** |
| | CoNi | $0.59 \pm 0.13$ | $0.78 \pm 0.06$ | 0.82 | *** |
| | CoZn | $0.71 \pm 0.05$ | $0.83 \pm 0.04$ | -0.02 | (n.s.) |
| | CuMn | $-0.15 \pm 0.10$ | $-0.21 \pm 0.05$ | 3.25 | *** |
| | CuNi | $0.90 \pm 0.02$ | $0.99 \pm 0.01$ | -0.98 | * |
| | CuZn | $0.87 \pm 0.02$ | $0.98 \pm 0.01$ | -1.76 | *** |
| | MnNi | $0.32 \pm 0.07$ | $0.36 \pm 0.07$ | 2.24 | *** |
| | MnZn | $0.02 \pm 0.04$ | $0.05 \pm 0.01$ | 2.68 | *** |
| | NiZn | $0.72 \pm 0.02$ | $0.81 \pm 0.06$ | 1.05 | *** |

† Mean ± SE over concentration levels 1-4
‡ $p(\chi^2; a = 0)$ given by asterisks: \* $< 0.05$ , \*\* $< 0.01$ , \*\*\* $< 0.001$

Combinations with copper tended to have the highest *δ* values, and those with manganese tended to have the lowest. Ranking the 15 combinations by *δ* (from low to high), the average rank of Cu combinations was 10.6, compared to an average rank of 3 for Mn combinations. Interestingly, the combination of copper and manganese displayed the lowest – and only negative – *δ* value.

### Quantifying deviations from concentration addition

For each metal combination, likelihood ratio tests were performed to determine whether concentration addition was rejected in favour of non-additive interactions (H0: *a* = 0, from equation 3). In Figure 2A we present examples of 3D graphical representations of the dose-response surface fitted by determining the maximum likelihood value of *a*. Notice that combinations with negative *a* values (synergistic interactions in CoCu and CdCu) have a depressed appearance in the center of the graph where fitness is lower when both metals are present compared to metals in isolation (fitness along the axes). The fitted surfaces of metal combinations that follow concentration addition (e.g., CdCo and CdZn; *a* not significantly different from 0) have a comparatively rounded appearance (e.g., considering the curve describing the EC50). Those with positive *a* values (antagonistic interactions in CoMn and CuMn) have dose-response surfaces with an extended ridge from the origin to the center of the graph, with higher fitness when both metals are present compared to either one in isolation. The maximum likelihood values of *a* ranged from −2.41 (“synergistic” with deep troughs) to +3.25 (“antagonistic” with shallow peaks), with the majority of combinations exhibiting antagonistic interactions (mean = 0.49, SD = 1.68; Figure 2B).

**Figure 2:**
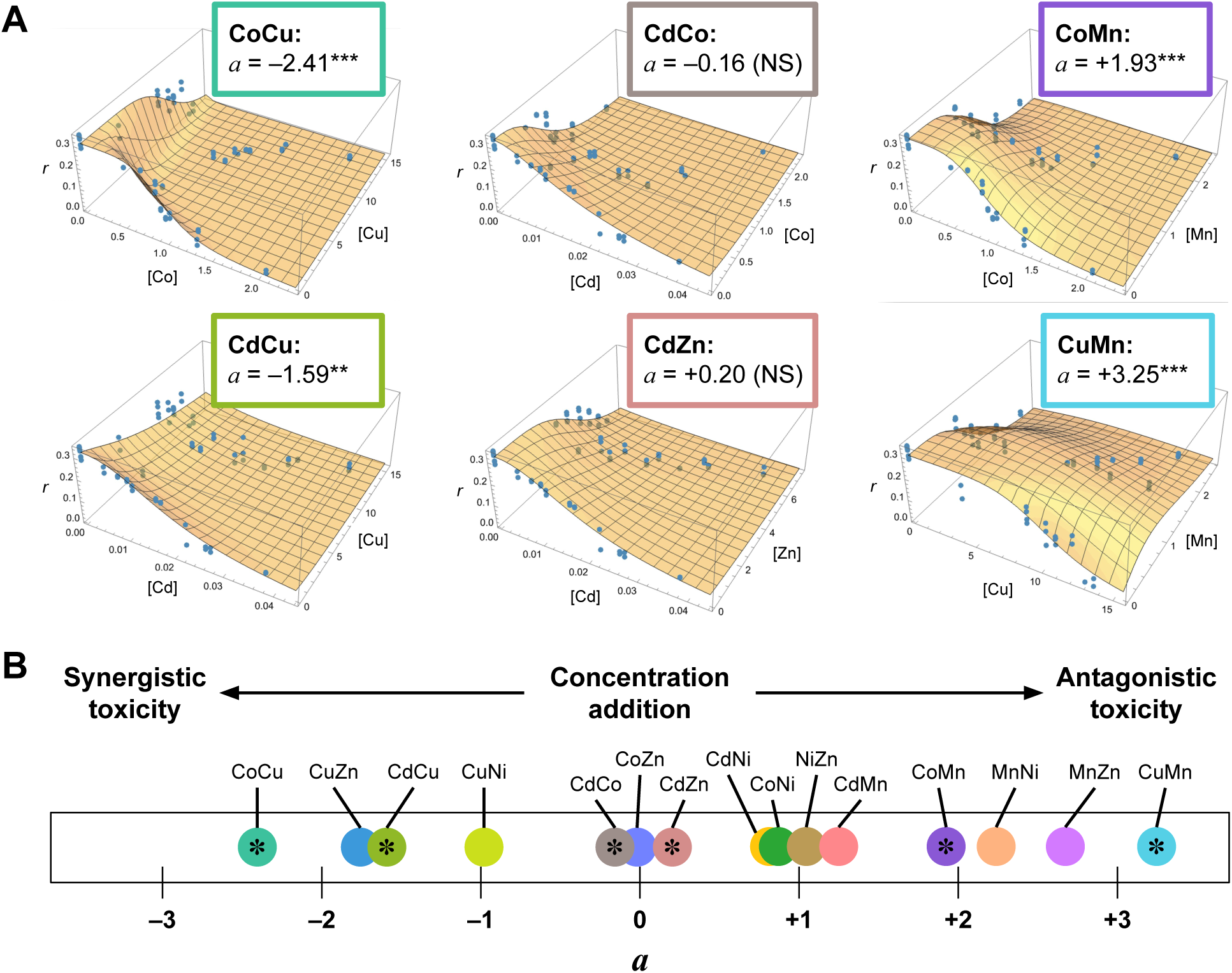
Metal interactions as estimated from the concentration-addition model. (A) Six examples of fitted surfaces with the metal combination and *a* values above each graph: *a <* 0 on left, *a* not significantly different from 0 in middle, and *a >* 0 on right. Significance level refers to improvement of fit when allowing for interaction (*a* ≠ 0 ***: *p <* 0.001). The vertical axis represents maximum growth rate (*r*); the horizontal axes represent concentration of each single metal (mM). See Figure S7 for all combinations. (B) Number line displaying *a* values and how they relate to synergism/antagonism. Dots with asterisks represent metal combinations displayed in panel A.

Three metal combinations interacted in a manner consistent with concentration addition (*a* = 0): CdCo, CdZn, and CoZn. Four combinations (CdCu, CoCu, CuNi, CuZn) exhibited significant synergistic toxicity (Figure 2B, Figure S7), with the *δ* values for the last three of these combinations near one, with negligible growth observed across all concentration levels tested (Figure 1). Notably, all four of these synergistically acting metal combinations included copper. By contrast, eight combinations exhibited significant antagonistic toxicity (Figure 2B, Figure S7), including the combinations with *δ* values near 0, which grew remarkably well across all concentration levels tested. Among these, manganese and nickel were over-represented, including all five combinations involving manganese and four of five combinations with nickel. Cadmium, cobalt, and zinc behaved neutrally in combination, with the value of *a* being determined by the other metal in the pair; that is, their combinations with copper were synergistic, their combinations with manganese or nickel were antagonistic, and their combinations with each other follow concentration addition (Figure 3).

**Figure 3:**
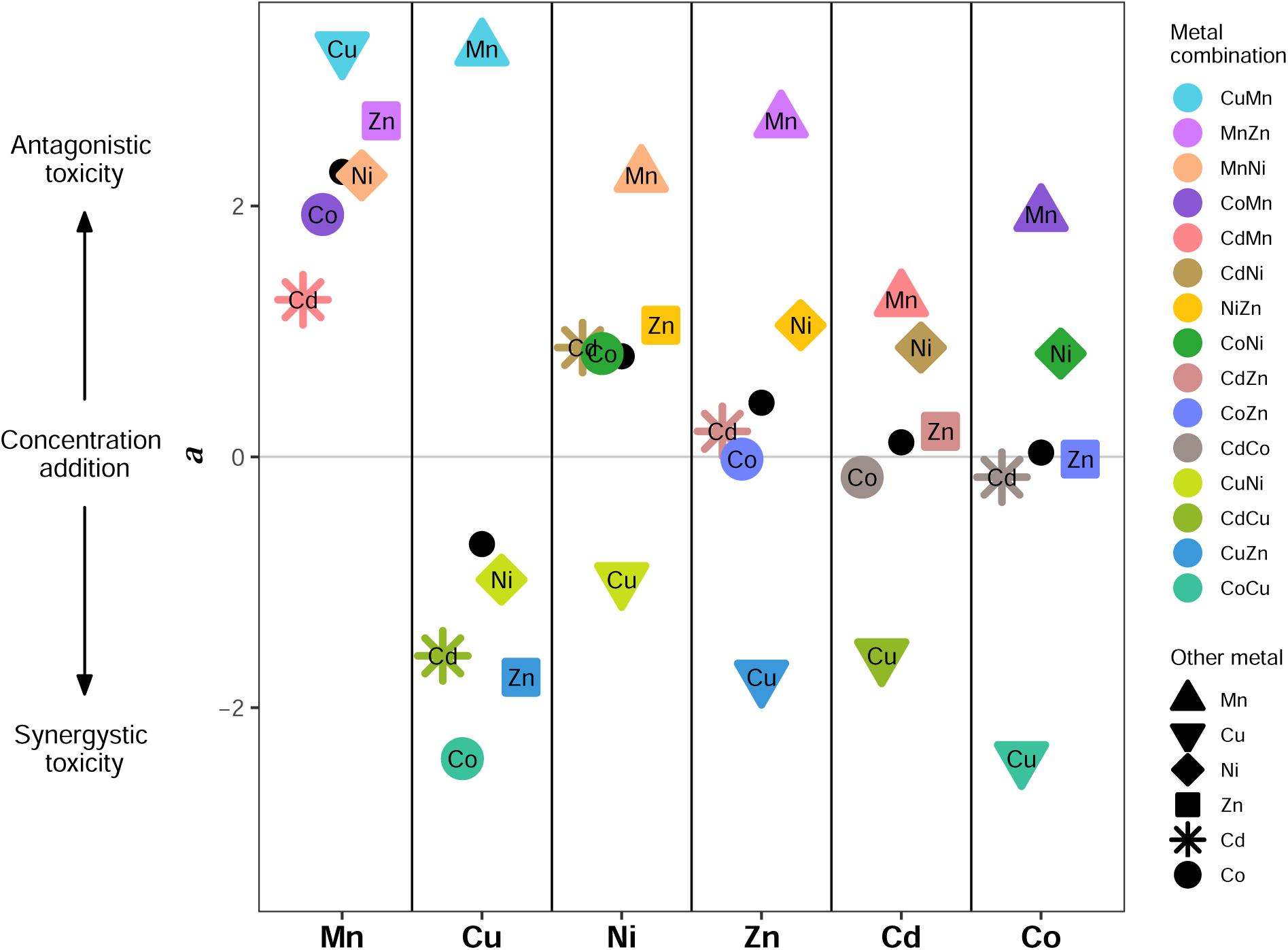
Interaction effects of individual metals. Single metals are shown along the x-axis, with their combinations displayed as colored points. The interaction value *a* is shown on the y-axis. Large black points are the average *a* value across combinations containing that metal.

The distribution of interactions experienced by each individual metal is displayed in Figure 3. The order of interaction strength among metals can be observed in the magnitude of *a* for Mn, Cu, and Ni’s combinations with the neutrally acting metals Cd, Co, and Zn (small open circles in Figure 3). The observed order of interaction strength is, therefore, Mn *>* Cu *>* Ni *>* Cd = Co = Zn. It is particularly striking that combining the most synergistic and antagonistic metals (i.e., Cu and Mn) results in the most antagonistic interaction, with growth nearly equal when combined as in the single metals (Figure 1A). We describe attributes of these metals that may account for these strong, but variable, interactions in the Discussion.

In Figure S8, we present similar results but calculating *a* from the difference in initial and final OD, rather than from maximum growth rates. There was a significant positive relationship between *a* calculated using growth rates and OD with a slope of of 0.91 (*R*^2^ = 0.63, *F* (1, 13) = 24.53, *p <* 0.001). This provides evidence that *a* is a fairly robust measure. References to *a* will subsequently refer to *a* as calculated from maximum growth rates.

### Comparison of *δ* and *a*

The two measures describing how stressors interact, *a* and *δ*, capture different attributes of mixture toxicity. *a* describes the deviation from concentration addition resulting from a chemical interaction between the two metals, with synergistic pairs having *a <* 0 and antagonistic pairs having *a >* 0. *δ*, on the other hand, describes the loss in fitness resulting from combining two metals compared to the average impact of the single metals. Statistically, *a* requires fitness estimates across a range of concentrations for each toxin to allow fitting the parameters in equations 2 and 3 to the dose-response surface, whereas *δ* requires only fitness with each of two single stressors of comparable magnitude and fitness in the combination. Nevertheless, the two measures are correlated with one another, and the concentration addition model provides a framework for understanding why *δ* remains roughly constant across a gradient of metal concentrations (Figure 4).

**Figure 4:**
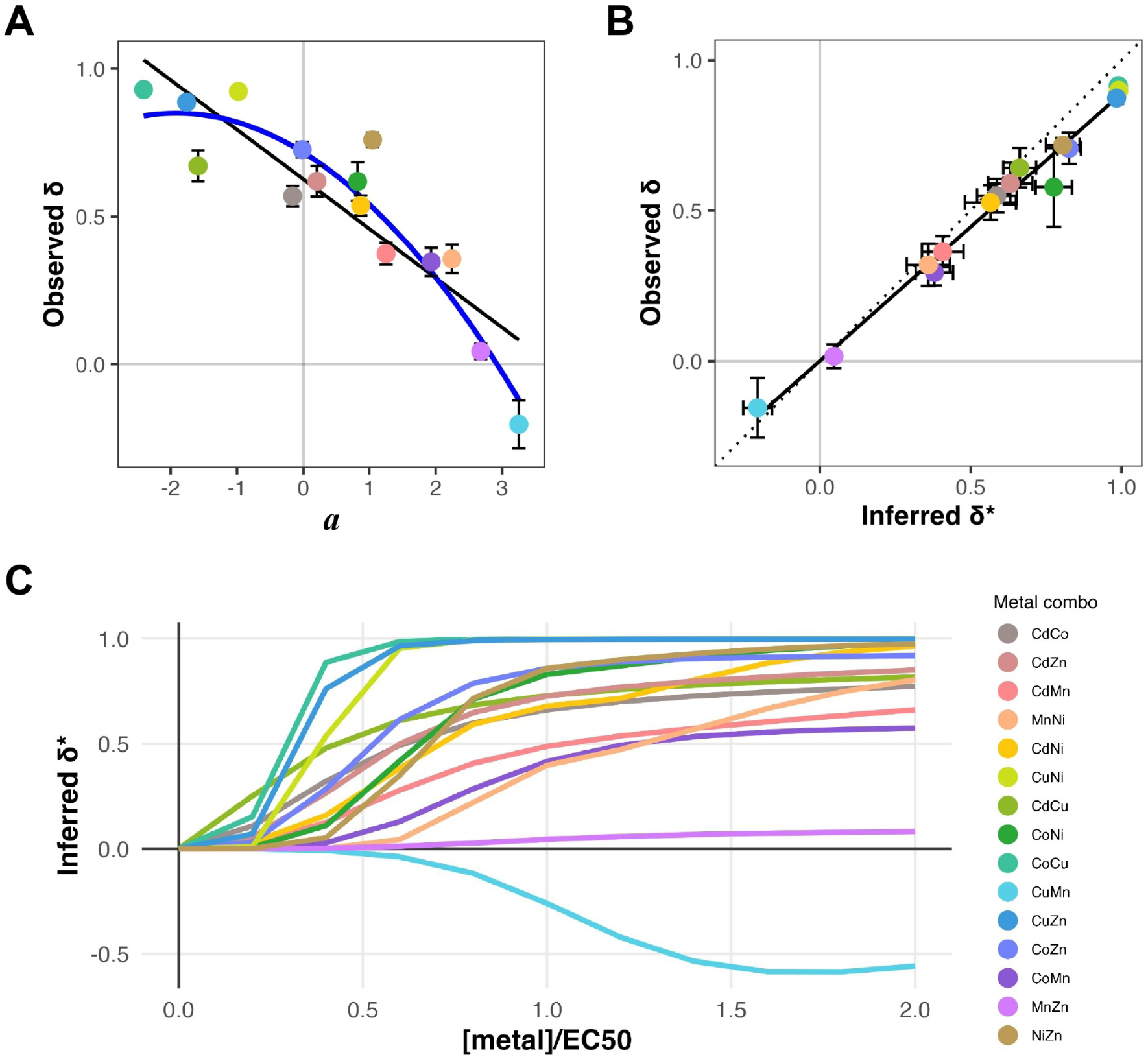
Relationship between *δ* and *a*. (A) Relationship between *δ* and *a*. (A) Relationship between the observed *δ* and the interaction value (*a*). The negative linear relationship (black line) is significant with a slope of (*R*^2^ = 0.76, *F* (1, 13) = 44.31, *p <* 0.001), but data fit even better to a quadratic function (blue line; *R*^2^ = 0.85, *F* (2, 12) = 41.94, *p <* 0.001) with AICs of −8.9 and −15.9, respectively. Error bars represent SE of the mean from *δ* measured at four concentrations. (B) Relationship between observed *δ* and model-estimated *δ*\*, where the latter is inferred from the dose-response surface using the maximum likelihood parameter estimates in equation 3 for each metal combination. The positive linear relationship is significant with a slope of 0.88 (*R*^2^ = 0.98, *F* (1, 13) = 704, *p <* 0.001). Dotted line marks a 1:1 relationship. Error bars represent SE of the mean from *δ* and *δ*\* measured at concentration levels 1-4. (C) Curves showing model-estimated *δ*\* values from the inferred dose-response surface for each metal combination, plotted across a range of stressor intensities. Color legend applies to all three panels.

Figure 4A shows the significant negative relationship between *δ* and *a* (*R*^2^ = 0.76, *F* (1,13) = 44.31, *p <* 0.001), a relationship that is even better fit by a second order polynomial curve with an accelerated decline (likely because *δ* cannot be larger than 1, at which point there is no growth in the combined metals). As expected, metals that when combined are very toxic (large *δ*) exhibit strongly negative synergistic interactions (negative *a*). Agreement between these metrics indicates that they measure the same underlying biological phenomenon, albeit using different methods.

The concentration-addition model also allows *δ* to be estimated at any metal concentration from the dose-response surface (e.g., Figure 2A) by simply determining the height of the surface for each single metal and their combination (inferred *δ*\* values are given in Table 2). That is, once parameters have been estimated, growth rates for any metal combination can be predicted from equation 3. We first did so at the five target metal concentrations and their combinations to obtain a model-estimated *δ*\* value. We find a strong linear relationship between the observed *δ* and estimated *δ*\*, with a slope of 0.89 (±0.07 95% CI) that is only marginally less than the 1:1 line (Figure 4B), indicating that these approaches are consistent with one another.

We also show in Figure 4C how *δ*\* is expected to change across stressor intensities, not just the five target concentrations. While *δ*\* values are near zero when metal concentrations are negligible, a striking feature of these model predictions is that *δ*\* should stabilize once the metals being combined approach or exceed their EC50 concentration (see also Figure S1 for similar plots across a range of *a* values). This provides a rationale for the use of *δ* as a good indicator of how harmful a metal combination is, as it is a fairly robust measure to the exact metal concentrations used. Interestingly, *δ*\* for CuMn does not stabilize until around 1.5 × EC50, consistent with the greater variation in observed *δ* across concentration levels (Figure 1C).

### Testing predictors of *δ* and *a*

Similarity between metals may contribute to their interactive dynamics when both are present. While simple physicochemical characteristics can describe similarities and differences of metals at an atomic level, it may be more relevant to consider metal similarity from the cell’s perspective. For example, if the cell uses the exact same cellular machinery for detoxification of two different metals, one might expect mutation (or altered regulation) of this machinery to effect resistance to both simultaneously. On the other hand, if the cell uses different structures for detoxification of each, conflict in resource allocation could dampen the cell’s ability to detoxify both metals at once. These scenarios consider similarity in cellular responses to metal toxicity, however, other metal characteristics may also predict similarity in how the cell experiences toxicity from each.

Bazzicalupo et al.^54^ present three similarity metrics for the predicting the evolution of cross-tolerance between metals in *Saccharomyces cerevisiae*: (i) oxidizing/reducing potential of media containing toxic concentrations of each metal, (ii) overlap in the effect of known metal resistance genes on the Saccharomyces Genome Database (SGD), and (iii) co-occurrence of metals in soil and sediment samples from the US Geological Survey (USGS). While Bazzicalupo et al.^54^ found that these metrics did not predict evolved crosstolerance, we anticipate they might have greater relevance for metal mixture toxicity. Therefore, we explored whether the interaction terms (*a* or *δ*) determined in this study showed a consistent relationship with these three aspects of metal similarity.

Relationships between the mixture toxicity metrics and the three predictors are displayed in Figure 5. Linear regression analysis reveals two significant patterns; there is a marginally significant positive relationship between *a* and the proportion of SGD resistance genes with the same effect (*R*^2^ = 0.21, *F* (1,13) = 4.66, p = 0.05). Thus, metal combinations with a higher proportion of overlapping genes affecting tolerance have milder interactions (antagonistic toxicity), potentially reflecting an evolutionary history of adaptation to these metal combinations. By contrast, there is a significant negative relationship between *a* and the difference in redox potential of single metals in yeast growth media (*R*^2^ = 0.28, *F* (1,13) = 6.53, p = 0.02). This result suggests that metals with very different oxidative properties combine in more harmful ways, potentially by compounding different stressors. Similar, but opposite, slopes were observed for *δ*, as expected from their relationship, although no other slopes were significant.

**Figure 5:**
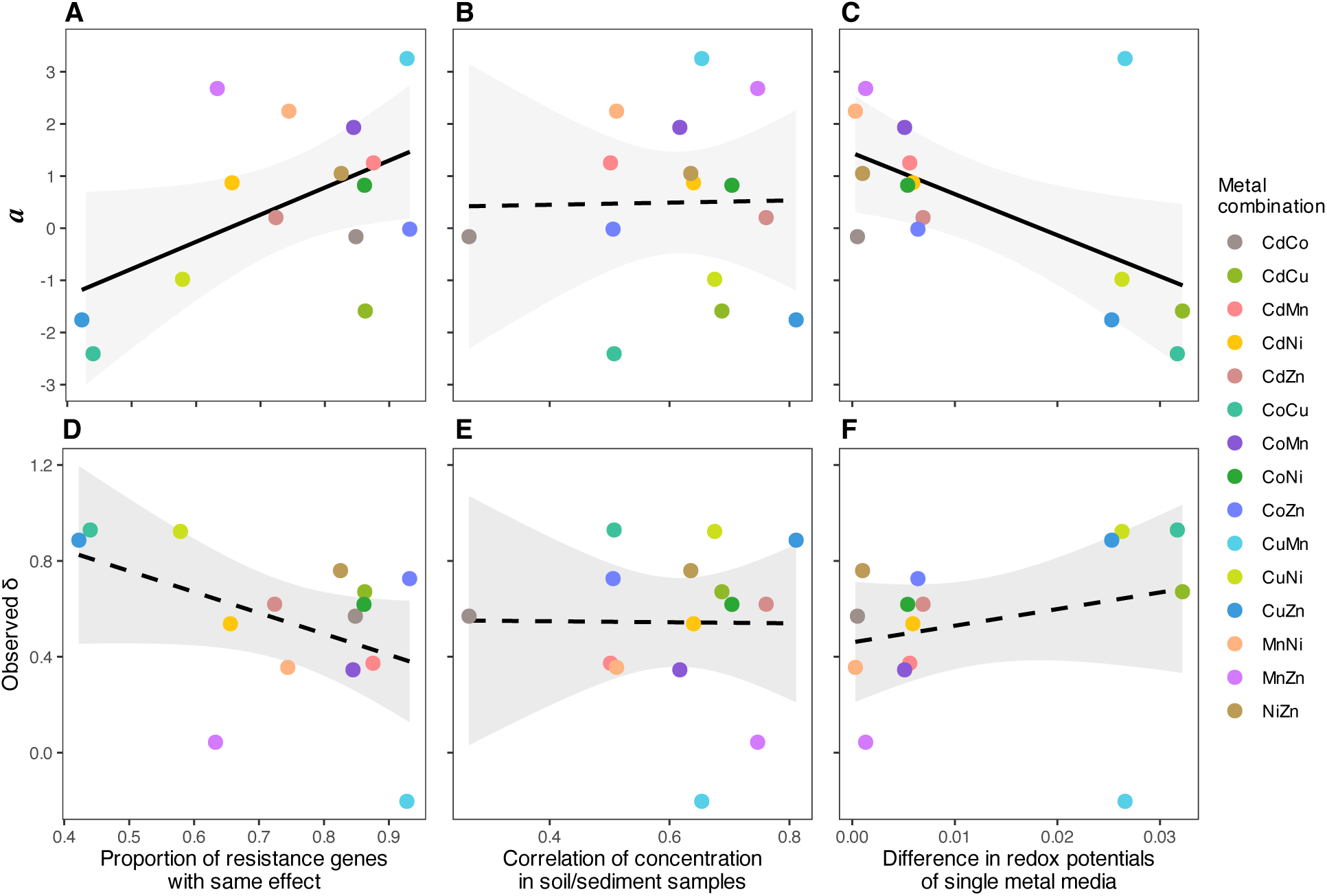
Predictors of metal interactions. The interaction terms for each combination (*a* in top row, *δ* in bottom row) are plotted against three measures of metal similarity (original analysis by Bazzicalupo et al. ^54^). (A) Linear regression of *a* and the proportion of metal resistance genes on SGD with the same effect (marginally significant; *R*^2^ = 0.21, *F* (1,13) = 4.66, *p* = 0.05). (B) Linear regression of *a* and the co-occurrence of metals in soil samples from the US Geological Survey (not significant). (C) Linear regression of *a* and difference in redox potential measured from single metal media (significant; *R*^2^ = 0.28, *F* (1,13) = 6.53, *p* = 0.02). Panels (D) through (F) show the relationship between *δ* and these same x-axes; none of these slopes are significant.

## Discussion

Both natural and human-induced stressors often co-occur, yet many ecological and evolutionary studies focus on the impacts on fitness of each stressor in isolation. Increasingly, there is recognition of the need to understand complex environmental changes^59^. The literature on chemical toxicity provides a foundation for measuring the impact of combining stressors^7^. Using *Saccharomyces cerevisiae* as a model, we have built on this foundation to assess which combinations of metals are particularly toxic when combined, posing a greater risk to natural populations.

We found considerable variability in the effects of metal combinations on yeast growth. To quantify how toxic combining metals is, we introduced a new measure, *δ*, that describes the difference in fitness when only one metal is present vs. when two equitoxic metals are combined (Figure 1B). A positive *δ* indicates decreased growth in the mixture, and the magnitude reflects the intensity of this effect. A *δ* value of 1 indicates maximum toxicity, where no growth is detected in the mixture regardless of growth in the individual metals. Metal combinations in this category include CoCu, CuNi, and CuZn (Figure 1C), indicating that these combinations may pose a much greater environmental hazard than expected from the single metals. For most metal combinations, exposure to two metals is more harmful than only one, as *δ* fell between 0.2 and 0.8, indicating a 20-80% reduction in growth relative to the single metals. Interestingly, there were two exceptions: combining MnZn led to virtually the same fitness as seen in each metal (*δ* of 0), and CuMn even showed a growth advantage in the mixture compared to the single metals (*δ <* 0).

We also described how the metals interact using the deviation from concentration addition, a null model that predicts that combining equitoxic amounts of two compounds should produce the same effect as doubling one of them^10,9^. Among the fifteen pairwise combinations of six metals (i.e., Cd, Co, Cu, Mn, Ni, Zn), three combinations behaved in a manner consistent with concentration addition (*a* = 0: CdCo, CdZn, CoZn). Four combinations displayed significant synergistic toxicity with harsher effects when combined (*a <* 0: CdCu, CoCu, CuNi, CuZn), and eight combinations displayed significant antagonistic toxicity with milder effects when combined (*a >* 0: CdMn, CdNi, CoMn, CoNi, CuMn, MnNi, MnZn, NiZn).

The harshest and synergistically toxic combinations – those with a negative *a* and very high *δ* values – all include copper. A possible contributing factor for the synergistic effect of copper with other metals is that copper is the most redox active of the metals tested here^60^. Copper ions readily participate in reactions that produce ROS (e.g., Fenton reactions, Haber-Weiss reactions), and ROS production is accepted to be the main mechanism of copper toxicity in many organisms^61,62^. Here, oxidative stress generated by these reactions may overwhelm the cell’s abilities to combat the toxic effects of the other metal. Furthermore, copper has been shown to permeabilize the plasma membrane to a greater extent than other metals, even at sub-lethal concentrations^63^, which may allow for greater influx of both metals into the cell.

The mildest and antagonistically toxic combinations – those with the highest *a* and low *δ* values – all include manganese. The antagonistic toxicity of these combinations may, in large part, be due to the antioxidant capabilities of manganese. Mn^2+^ plays a critical role as a cofactor in the superoxide dismutase MnSOD, which converts deadly superoxide to oxygen or hydrogen peroxide^64^. MnSOD is usually located in the mitochondrion but can function in the cytoplasm under manganese-replete conditions^65^. Mn^2+^ is also known to form complexes with small molecules (e.g., phosphates, carbonates), which function as effective antioxidants, scavenging harmful ROS from the cytoplasm and environment^66^. Having an abundance of these manganese antioxidants may prevent oxidative damage in the cell, despite a large volume of ROS produced by the other metal in the environment. It is possible that the manganese’s antagonistic effect on the toxicity of the other metal is strongest for metals whose main mechanism of toxicity is ROS production. For example, the antagonistic effect of manganese is strongest with copper (most active in ROS production) and weakest with cadmium (least active in ROS production)^67^.

Finally, metal combinations that do not display strong interactions consist of cadmium, cobalt, and zinc. These metals appear to play a more neutral role when combined with other metals, such that the sign of the interaction term (*a*) depends on the other metal with which they are paired.

One major mechanism by which excess metal concentrations reduce fitness is by replacing natural cofactors in enzymes. Briefly, the binding affinity of a divalent metal ion for a given ligand is correlated with its second ionization energy (i.e., the energy required to remove a second electron from a 1+ ion)^15^. Based on this principle, the Irving-Williams series ranks the relative stability of metal-ligand complexes in the following order: Mg^2+^ *<* Mn^2+^ *<* Fe^2+^ *<* Co^2+^ *<* Ni2+ *<* Cu^2+^ *>* Zn^2+^ (zinc is traditionally placed to the right of copper to preserve the order of atomic weights in the periodic table)^68^. Copper’s high binding affinity makes it a strong competitor to replace native cofactors in enzymes, and once inside, it may cause oxidative damage to the protein. By contrast, manganese is not a strong competitor for these binding sites, and therefore is unlikely to displace native cofactors. Our findings indicate that combining metals that are both high in the Irving-Williams series are particularly toxic (Figure 1).

Literature on the mechanisms of nickel toxicity is relatively scant, so, although the following speculations provide a possible explanation for nickel’s antagonistic effects, they can be regarded with reservation. Nickel is relatively flexible in assuming various coordination geometries. Coordination geometry refers to the orientation of ligands bound to a metal ion, such as the amino acids surrounding a cofactor, and depends on ion size and electron configuration^69^. Compared to other metals, nickel can adopt any of the most common coordination geometries (i.e., octahedral, tetrahedral, square planar, and linear) with greater thermodynamic stability^70^. An enzyme that has had its native cofactor competitively replaced is more likely to retain function if the competitive ion is able to adopt the coordination geometry of the native cofactor^15^. Given that nickel may be more flexible in its coordination geometry, it may be better able to preserve enzyme function when replacing native cofactors. In addition, because it has a relatively high binding affinity^68^, nickel may reduce the harm of metal combinations because it outcompetes other ions that would cause loss of enzymatic function due to changed confirmation of the protein.

It may be the case that the true mechanisms underlying the interactions we observed are not captured by our review of the literature. For example, metals also impact membrane transport, vacuolar storage, and transcriptional regulation, among other cellular processes, each of which could vary nonadditively in a metal mixture. Additionally, the general stress response of a cell may exhibit non-linear responses in the face of multiple stressors. These complex physiological processes, as well as metal-specific uptake kinetics and intracellular compartmentalization, likely also contribute to the diverse interaction patterns we observed. Additionally, the growth medium used as the background environment for this experiment is composed of a highly complex mixture of organic molecules, which may also interact with metal ions, differentially impacting their bioavailability and reactivity^71^.

Interestingly, we found that metal combinations that have more genetic overlap in known metal resistance genes in yeast tend to have higher *a* values (i.e., milder antagonistic interactions), potentially due to an evolved tolerance to these combinations. By contrast, combinations of metals with a greater difference in redox potential had more negative *a* values (i.e., harsher synergistic interactions).

While *a* and *δ* are both measures of mixture toxicity, they capture different aspects of metal interactions. The parameter *a* quantifies deviations from concentration addition by modifying the curvature of a fitted 3D dose-response surface, providing a model-based assessment of synergy and antagonism across a gradient of metal concentrations. In contrast, *δ* is a relatively simple, experimentally direct measure that compares growth in the mixture to the average growth in the individual metals. Despite its simplicity, *δ* is not merely a snapshot metric; our analysis shows that it remains relatively consistent across concentrations for a given metal combination (Figure 4C) and is significantly correlated with *a* (Figure 4A), indicating that it captures intrinsic properties of metal-metal interactions. Thus, while *a* provides a theoretical framework for evaluating mixture toxicity, *δ* offers an intuitive and broadly applicable metric of the impact of mixing equitoxic stressors that aligns with these modeled interactions.

Furthermore, the difference in toxicity of combinations compared to individual components (*δ*) may often be more relevant for ecotoxicological applications than overall synergism or antagonism, as it directly reflects organismal growth and survival. While synergism and antagonism describe how toxicity deviates from an expected additive model, the absolute reduction in fitness caused by a mixture of toxins may provide a more practical measure of ecological risk, particularly in scenarios where it is necessary compare the magnitudes of effect between different mixtures.

Empirical evidence of metal mixture toxicity from other studies suggests that patterns of synergism and antagonism between metals are sensitive to study species and methodology used to assess the interaction. For example, while our data using yeast growth rate shows that CdCu and CuZn are synergistic and CdZn follows CA, analyses from fifteen other studies using various methodologies and study organisms show all possible interaction types for these metal combinations (Table S2). The measure of effect for determining the interaction between metals can vary according to the natural history of the organism being studied (e.g., mortality/survival in animals, root growth/inhibition in plants, population growth rate or fluorescence in microbes, etc.), and this variation may cause the conclusions from each study to differ. Furthermore, evolution can shape these interactions^56^ by altering the structure and function of relevant cellular machinery. Despite the limited number of studies and metal combinations, there is a significant positive relationship between reported interactions and *a* as measured in our study (Figure S9), using a qualitative score for reported interactions (from −1 for synergism to +1 for antagonism).

Notably, studies of fungi leave a large gap in the metal mixture literature. Mesquita et al.^72^ represents the only fungal investigation of metal mixture toxicity for cadmium, lead, and zinc, but their focus was on biochemical indicators of toxicity (e.g., ROS accumulation, membrane integrity) rather than quantifying the effect on fitness. Most of the combined metal research in fungi is focused on uptake, biosorption, and bioremediation rather than toxicity^73,74,75,76,77,78^. It is possible that fungal cell features, such as the chitinous cell wall and fungal-type vacuole, contribute to differences in how metal interactions affect their structure and function. There is thus value to investigating the toxicity of metal combinations specifically in fungi. As fungi are vital components of natural ecosystems and can contribute to detoxification of polluted environments, understanding how fungi withstand complex environmental stresses is vital for their conservation and potentially for their use in bioremediation.

Further studies will be required to validate the generalizability of the *δ* measure for other stressor combinations and study organisms. Given its simplicity and direct applicability, *δ* could serve as a useful comparative metric across different taxa and environmental contexts, but further testing is needed to determine its robustness beyond metal toxicity and yeast population growth rates. Our work measured maximum growth rate of a microbial population as the response variable, but other taxa might require different measures that are more relevant to their biology or easier to measure in that system. For instance, fitness metrics such as survival rates, reproductive success, or physiological stress markers might be more appropriate in plants or animals, and assessing *δ* in these contexts could help determine its broader applicability. It may be the case that the *δ* framework holds its validity for dissimilarly acting stressors, like temperature and pH, but this remains to be seen. Future work should explore whether *δ* consistently captures interaction effects in these contexts or if modifications are needed to account for stressors with fundamentally different modes of action.

