## Supplemental Material for "Quantifying the effect of metal interactions on growth rate in *Saccharomyces cerevisiae*"

### Appendix

#### Supplementary Tables

Table S1: Experimental concentrations. Two independent fitness trials were conducted, targeting five concentrations of the single metals that would yield maximum growth rates of {0.25, 0.2, 0.15, 0.1, 0.05} (labeled as concentration levels 1 to 5, respectively). The second trial used slightly modified concentrations to better reach these targets, although the growth differences between trials were minor (Figure S3H)

| | | Concentration level (mM, below) | | | | |
|---|---|---|---|---|---|---|
| Metal | Trial | 1 | 2 | 3 | 4 | 5 |
| Cd | 1 | 0.008 | 0.011 | 0.017 | 0.026 | 0.038 |
| | 2 | 0.005 | 0.010 | 0.016 | 0.022 | 0.037 |
| Co | 1 | 0.70 | 0.86 | 1.00 | 1.32 | 2.05 |
| | 2 | 0.50 | 0.85 | 0.90 | 0.95 | 1.30 |
| Cu | 1 | 8.83 | 10.17 | 10.83 | 12.00 | 13.50 |
| | 2 | 4.00 | 8.83 | 10.17 | 10.85 | 12.90 |
| Mn | 1 | 0.86 | 1.14 | 1.31 | 1.66 | 2.23 |
| | 2 | 0.85 | 1.10 | 1.14 | 1.25 | 1.60 |
| Ni | 1 | 1.88 | 2.40 | 3.00 | 3.60 | 5.10 |
| | 2 | 2.30 | 2.60 | 2.75 | 3.00 | 3.20 |
| Zn | 1 | 4.00 | 4.75 | 5.40 | 6.00 | 6.60 |
| | 2 | 3.33 | 4.00 | 4.80 | 5.40 | 6.00 |

Table S2: Metal interactions from previous studies. Qualitative reports of metal interactions are shown for three commonly studied metal pairs. ANT: Antagonistic; SYN: Synergistic; ADD: Additivity.

| Author | Year | Organism (species) | Response measure | CuZn | CdCu | CdZn |
|---|---|---|---|---|---|---|
| Crémazy et al. | 2018 | Snail (*L. stagnalis*) | Individual growth rate | ANT | ANT | ANT |
| Franklin et al. | 2002 | Alga (*Chlorella* spp.) | Population growth rate | ANT | SYN | ANT |
| Gao et al. | 2018 | Zebrafish (*D. rerio*) | Survival | ADD | SYN | — |
| Gao et al. | 2020 | Cyanobacteria (*M. aeruginosa*) | Population growth rate | SYN | ANT | ADD |
| Gopalapillai & Hale | 2015 | Duckweed (*L. minor*) | Root growth | — | ANT | — |
| Liu et al. | 2015 | Lettuce (*L. sativa*) | Root growth | — | SYN | — |
| Lock & Janssen | 2002 | Potworm (*E. albidus*) | Reproduction | ANT | ADD | ADD/SYN |
| Lynch et al. | 2016 | Fathead minnow (*P. promelas*) | Mortality | SYN | — | — |
| Meyer et al. | 2015 | Daphnia (*D. magna*) | Mortality | SYN | ADD/SYN | ANT |
| Nagai & De Schamphelaere | 2016 | Diatom (*N. pelliculosa*) | Population growth rate | ANT | ANT | ANT |
| Otitoloju | 2002 | Snail (*T. fuscatus*) | Mortality | ADD | ANT | ANT |
| Preston et al. | 2002 | Bacteria (*E. coli*) | Luminescence | SYN | SYN | SYN |
| | | Bacteria (*P. aeruginosa*) | | ADD | ADD | ADD |
| Utgikar et al. | 2004 | Bacteria (*V. fischeri*) | Luminescence | SYN | — | — |
| Versieren et al. | 2007 | Barley (*H. vulgare*) | Root growth | — | ADD | ADD/ANT |
| Xu et al. | 2011 | Sea urchin (*S. intermedius*) | Abnormal developoment | ADD | ADD | ANT |

### Supplementary Figures

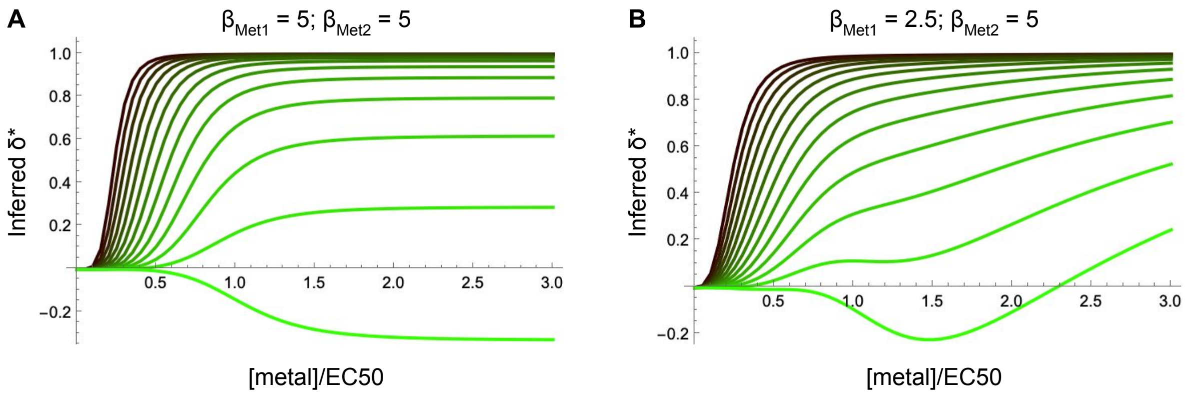


Figure S1: Curves show model-estimated $\delta^*$ values obtained from dose-response surfaces across a range of stressor intensities (x-axes). Color indicates the interaction value $a$, ranging from $-3$ (dark green; synergistic toxicity) to $+3$ (light green; antagonistic toxicity). Panels show different values of $\beta$ describing the steepness by which fitness declines in dose-response curves in the two metals: (A) $\beta[1]$=$\beta[2]$=5, (B) $\beta[1]$=2.5; $\beta[2]$=5. Figure 4C is similar but uses the parameters estimated from the data for each metal combination (i.e., $a$, $max$, EC50[1], EC50[2], $\beta[1]$, $\beta[2]$).

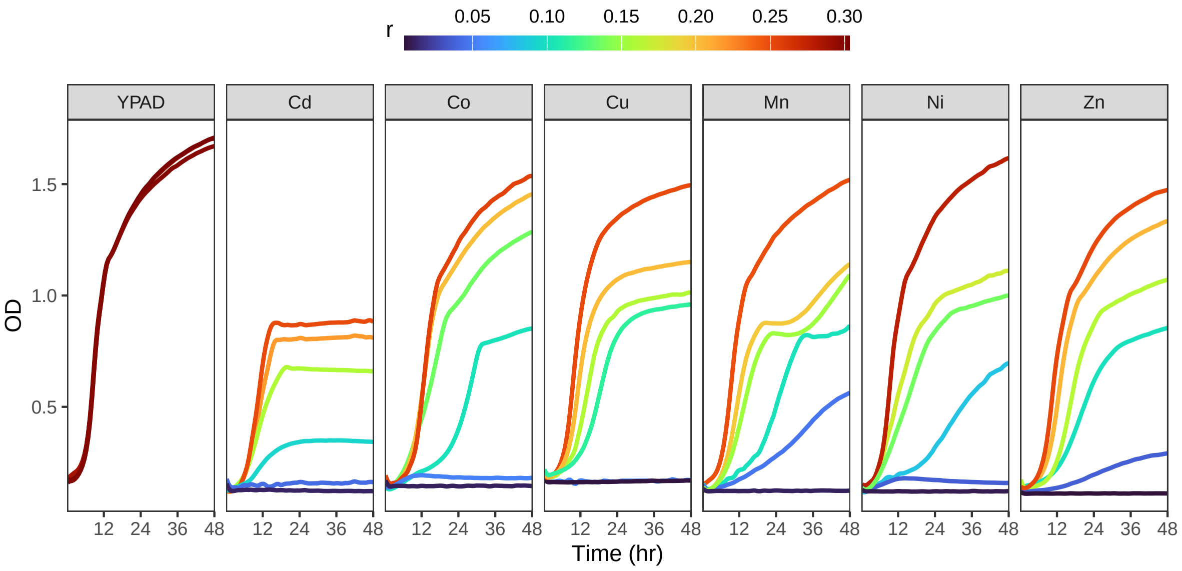


Figure S2: Example growth curve data of OLY077 measured in the Bioscreen C. These are examples of growth curves used to calculate maximum growth rates ($r$) displayed in Figure 1A). These curves are loess-fits to OD measurements using the same settings used in the calculation of $r$. The colour of the line shows the value of the maximum growth rate ($r$) calculated using the procedure described in the Methods. On the far left are typical growth curves of OLY077 in YPAD with $r \cong 0.3$. Each graph to the right shows examples of OLY077 growth curves at a range of concentrations levels of the metal identified at the top of the graph. The black lines, with $r$ near 0.00, are all blanks (containing no yeast) of the lowest concentration. Metal concentrations can be found in Table S1.

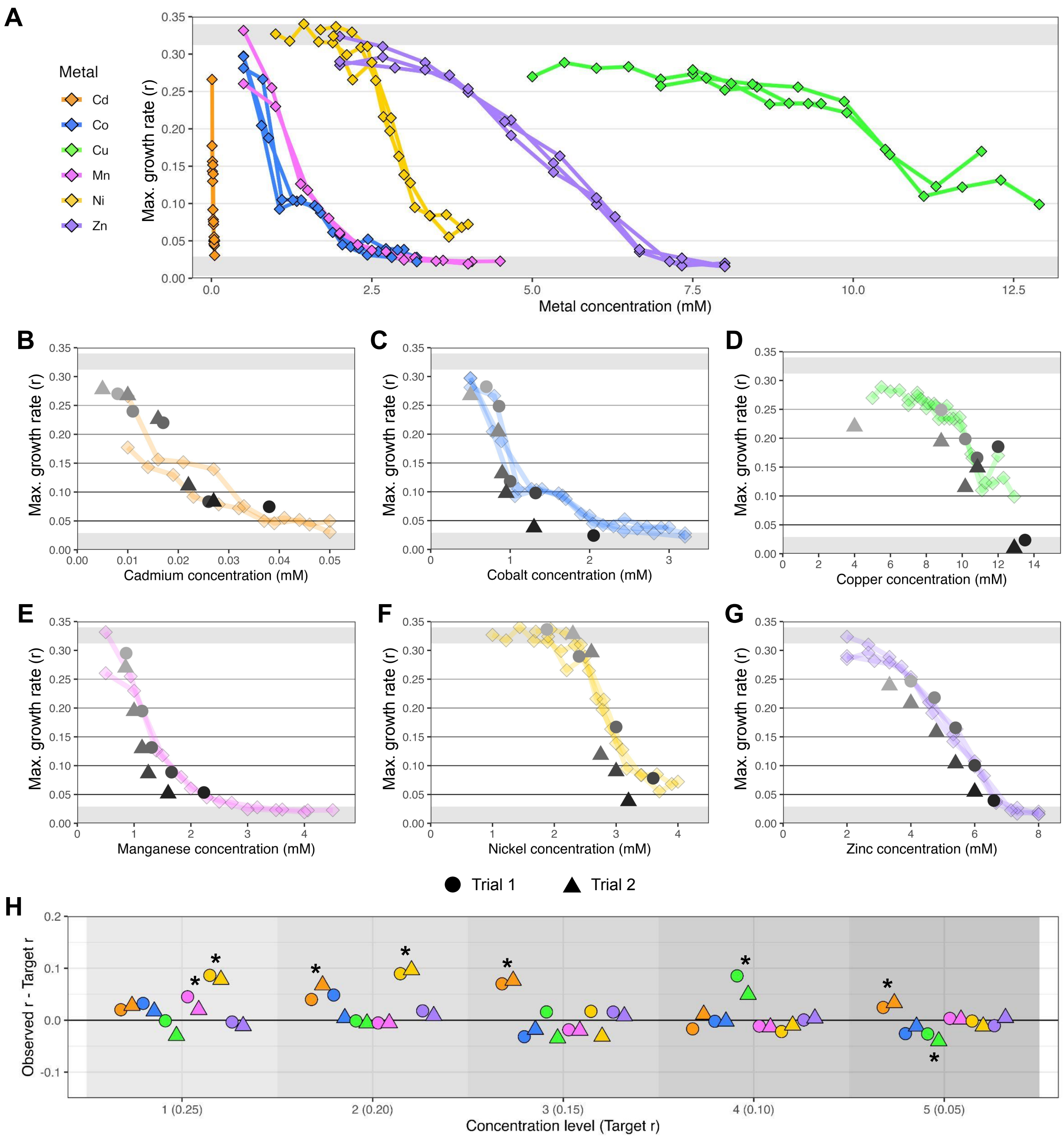


Figure S3: Dose-response curves for individual metals. (A) Results from initial experiments conducted with each single metal to determine their dose-response curves. The x-axis is the concentration of metal (mM), and the y-axis is the maximum growth rate ($r$). Color denotes metal; lines represent independent trails. Light grey bars on top and bottom display range of positive and negative controls, respectively (positive control, yeast + YPAD; negative control, water + YPAD). (B-G) Growth rates from these same single metals estimated in our experiment, along with the previously estimated dose-response data from panel A in color; grey value corresponds to concentration level and point shape to trial, as in Figure 1. Horizontal lines mark target $r$ for single metals in the combination assay. Note the y-scale is constant, but the x-scale is specific to metal. (B) Cadmium. (C) Cobalt. (D) Copper. (E) Manganese. (F) Nickel. (G) Zinc. (H) Difference in the observed and target growth rates for the five different target concentration levels. Point color represents metal as in panel A; point shape represents trial as in panel B. Groups along x-axis represent concentration level matching the target $r$ displayed as horizontal lines in panel B.

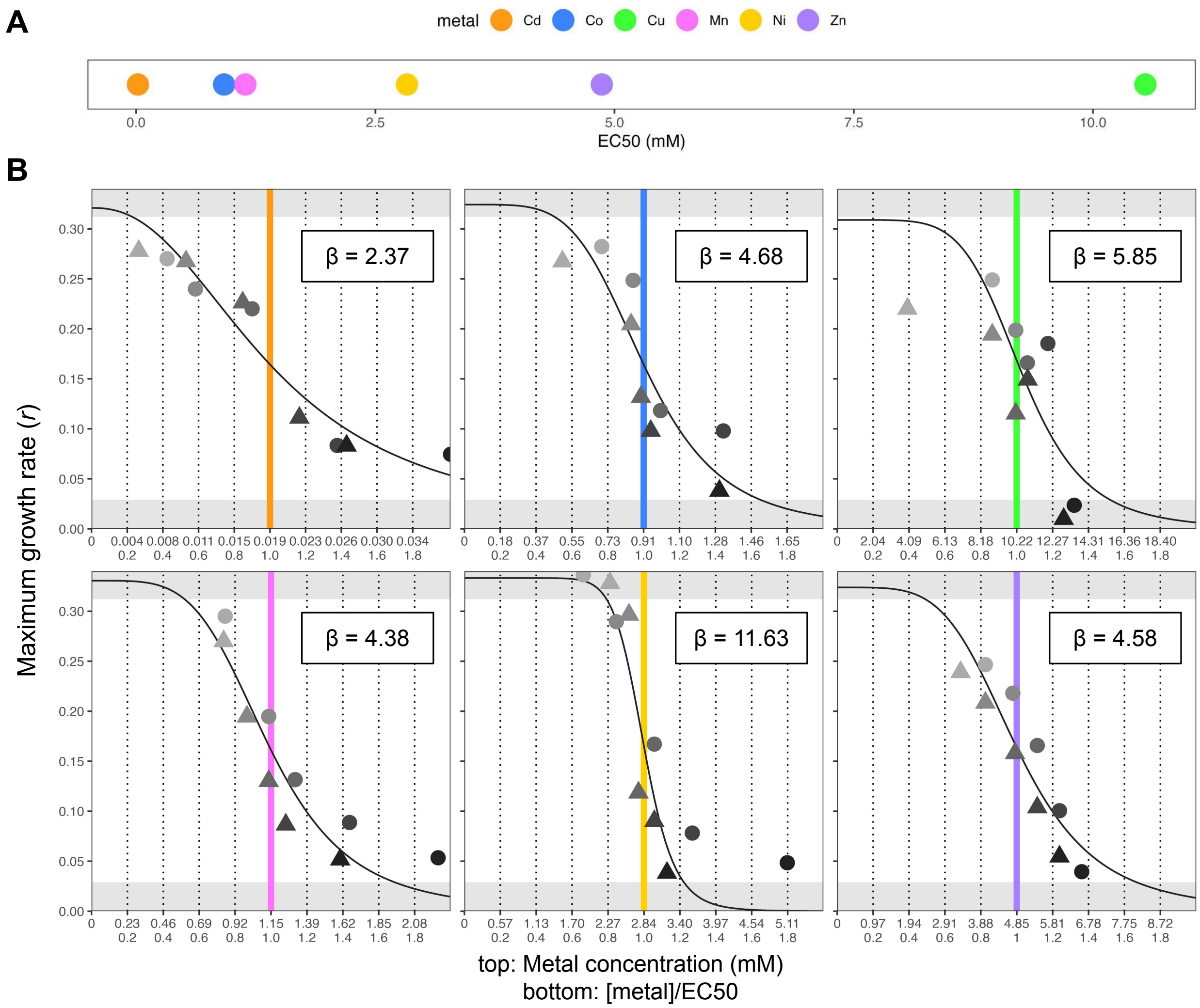


Figure S4: Demonstration of how $\beta$ affects dose-response curve. Equation 2 was used to estimate the maximum likelihood parameter values for $max$, EC50, $\beta$ given data on the maximum growth rates across in each single metal (points, showing the same data as in Figure S3).

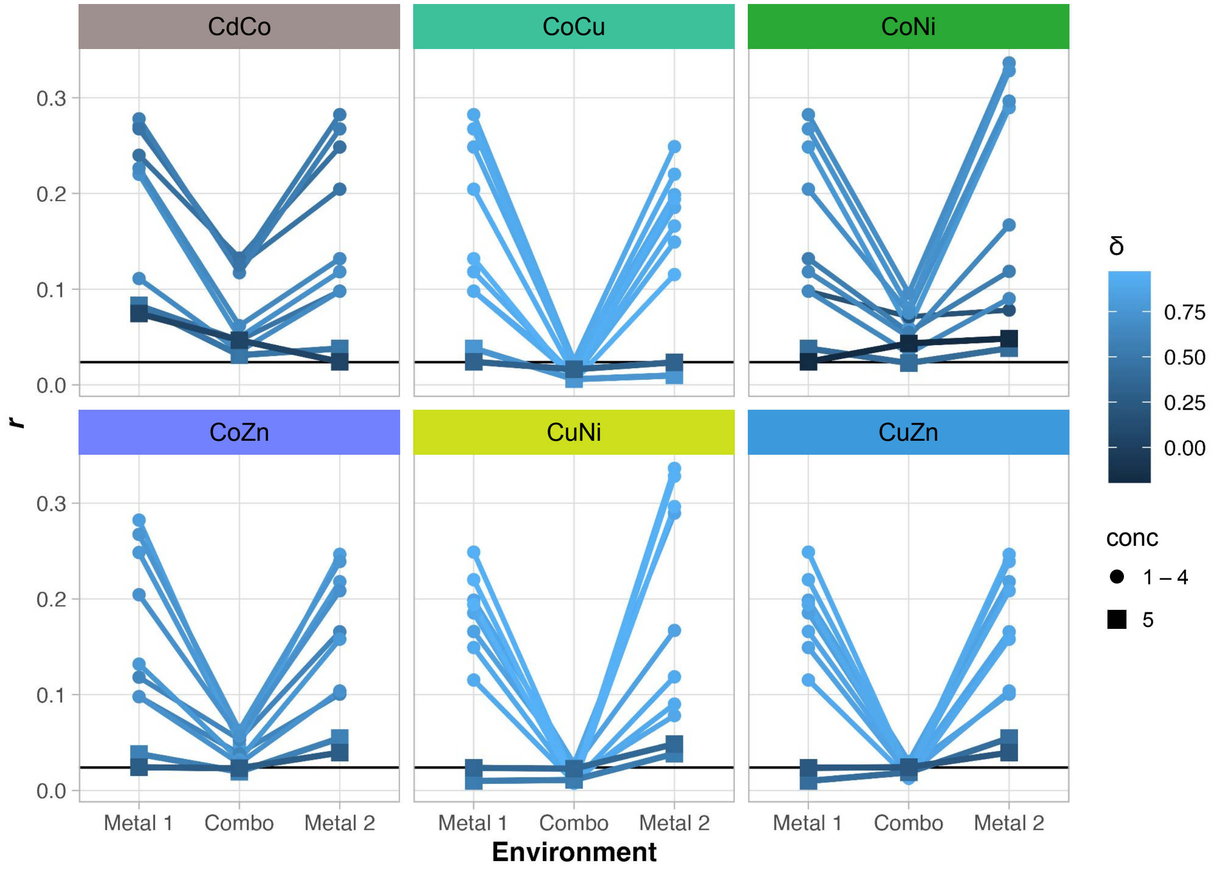


Figure S5: Low growth at concentration level 5. (A) Example graphs of yeast growth rates in 6 metal combinations (same data as in Figure 1A). Data represent two independent experiments. Lines connect samples in the same concentration level and trial, and color value represents $\delta$ calculated from each. Small circular points represent data from concentrations 1-4, and square points represent concentration 5. The latter estimates are so low that they are hard to distinguish from the negative controls lacking yeast, eliminating the signal needed to estimate $\delta$ accurately.

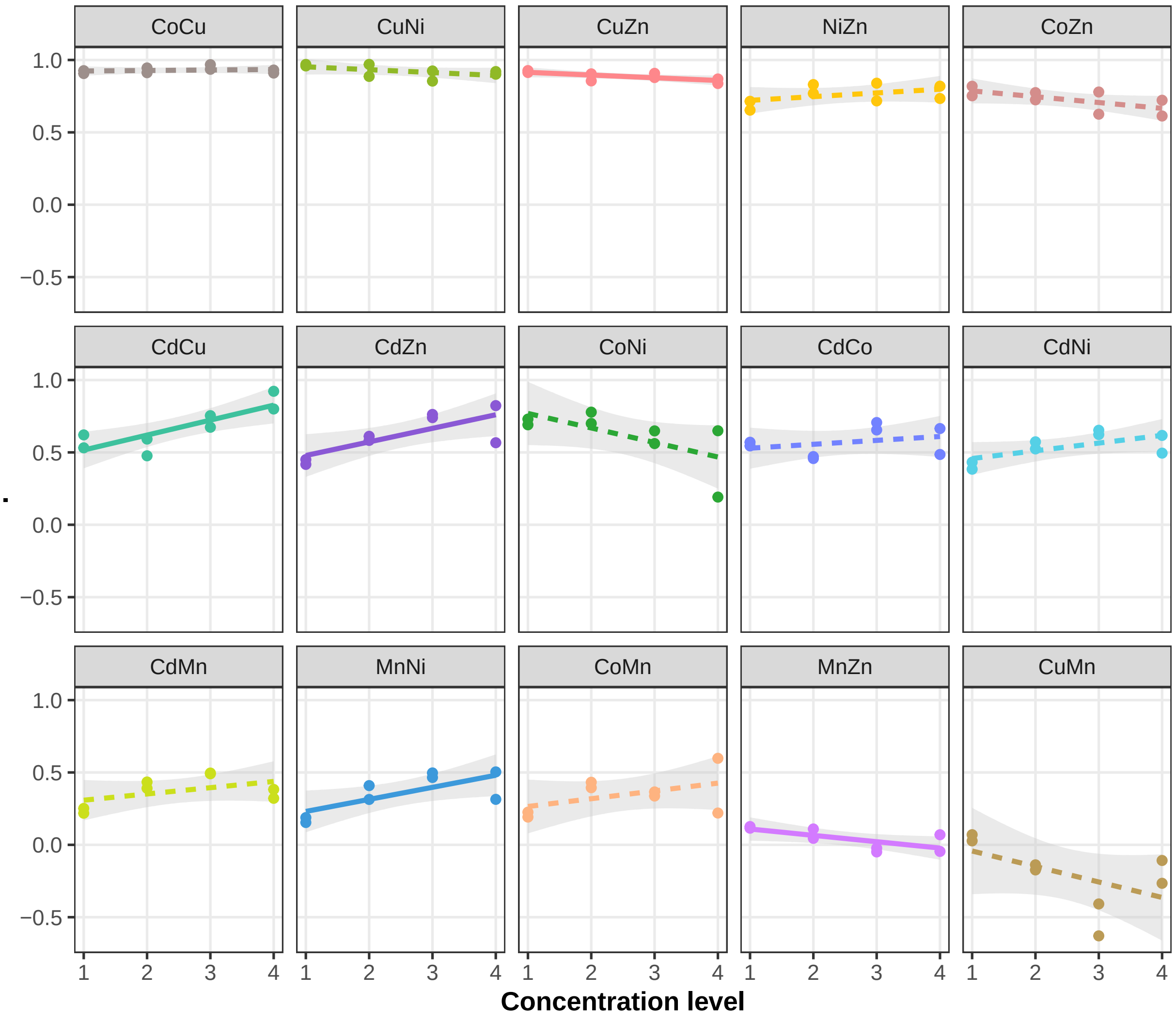


Figure S6: Relationship between $\delta$ and concentration level in each metal combination. Each graph represents a metal combination. Concentration level is given on the x-axis, and $\delta$ is given on the y-axis. Dotted lines represent correlation with $p > 0.05$, solid lines represent correlation with $p < 0.05$, though no correlations remain significant after Bonferroni correction for multiple comparisons. Grey area around the line represents the 95% confidence interval.

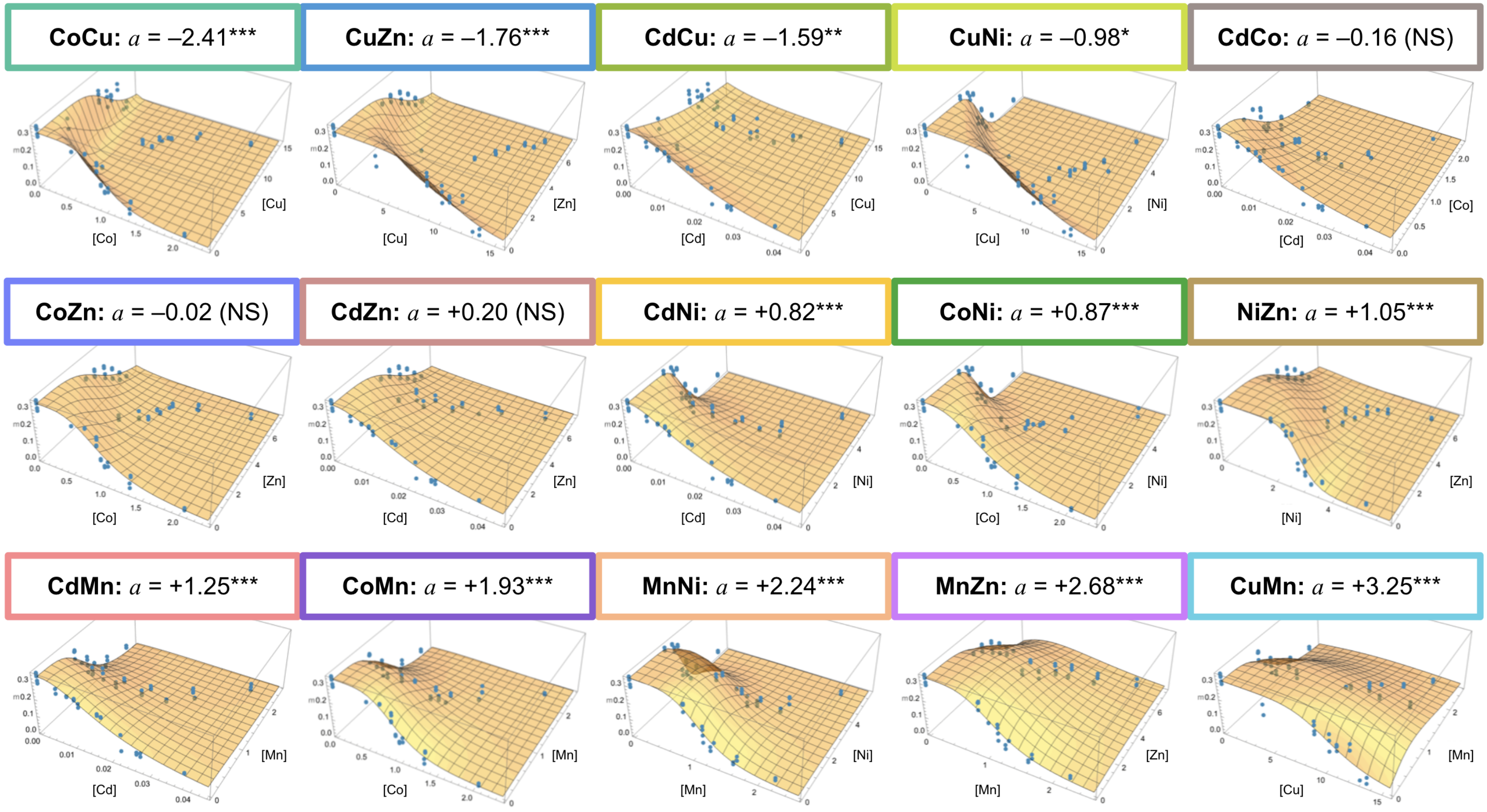


Figure S7: Fitted 3D surfaces and interaction values ($a$) from all pairwise combinations. Significance level refers to improvement of fit when allowing for interaction ($a \neq 0$ ***: $p < 0.001$). The vertical axis represents maximum growth rate ($r$); the horizontal axes represent concentration of each single metal (mM).

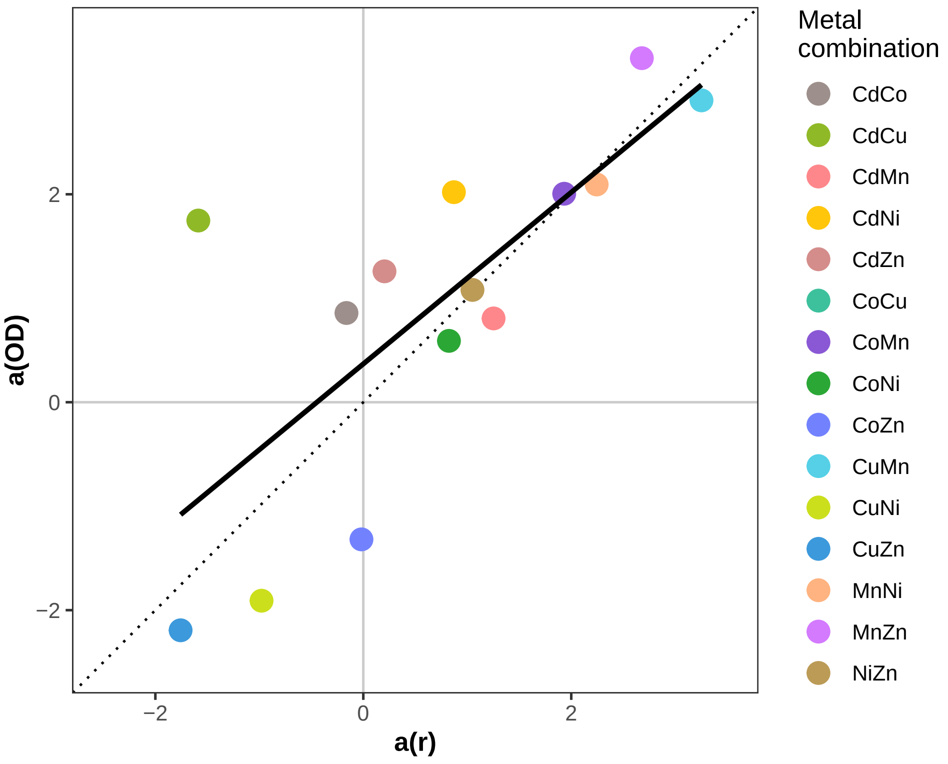


Figure S8: Relationship between $a(\mathrm{OD})$ and $a(r)$. There is a significant positive relationship in the $a$ value determined using the difference between final and initial OD (y-axis) and the $a$ value determined using maximum growth rates (x-axis) with a slope of 0.91 ($R^2 = 0.63$, $F(1, 13) = 24.53$, $p < 0.001$). The dotted line represents the 1:1 line.

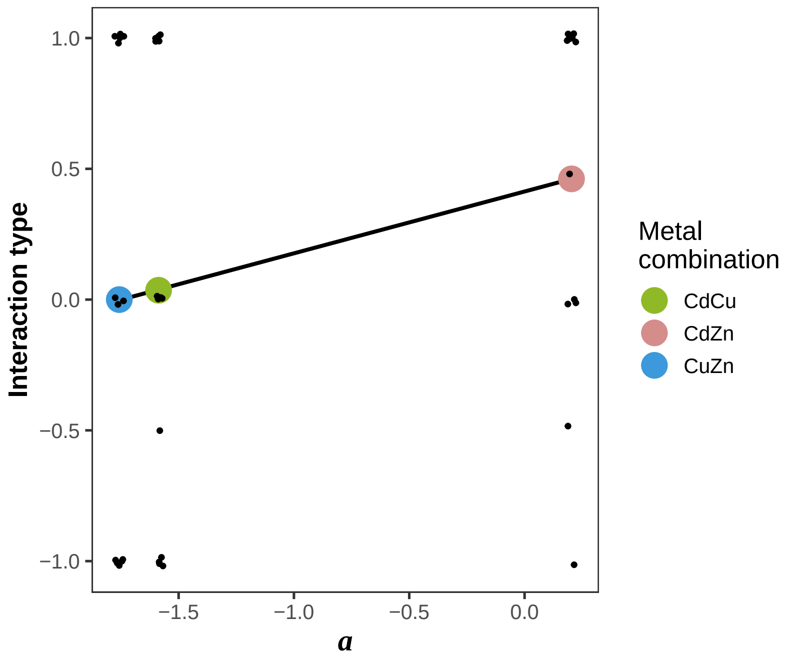


Figure S9: Relationship between metal interactions described in the literature and interaction term $a$ calculated in the present study. Results from fifteen papers assessing mixture toxicity in combinations of CdCu (n=14), CdZn (n=13), and CuZn (n=13) were compiled (see Table S2 for details). The value of $a$ as calculated in the present study is shown on the x-axis. As quantitative measures of interactions were generally not available, a qualitative assessment of interaction type is shown on the y-axis, where synergism receives a value of $-1$, additivity receives $a$ value of 0, and antagonism receives a value of $+1$. Some papers reported a combination of additivity/synergism or additivity/antagonism depending on conditions, resulting in values of $-0.5$ and $+0.5$, respectively. Small black dots represent results from different studies. Large colored dots represent the average interaction across studies. The linear relationship for average interaction versus a was significantly positive with a slope of 0.24 ($R^2 = 0.99$, $F(1, 1) = 1.2 \times 10^4$, $p < 0.01$).
